# Coxsackievirus B3 elicits a sex-specific CD8^+^ T cell response in female mice

**DOI:** 10.1101/2022.08.09.503381

**Authors:** Adeeba H. Dhalech, Stephanie A. Condotta, Caleb Corn, Martin J. Richer, Christopher M. Robinson

## Abstract

Sex is a significant contributor to the outcome of human infections. Males are frequently more susceptible to viral, bacterial, and fungal infections, which is often attributed to a weaker immune response. In contrast, a heightened immune response in females enables better pathogen elimination but leaves females more predisposed to autoimmune diseases. Unfortunately, the underlying basis for sex-specific immune responses remains poorly understood. Here, we show a sex-specific difference in the CD8^+^ T cell response to an enteric virus, Coxsackievirus B3 (CVB3). We found that CVB3 induced expansion of CD8^+^ T cells in female mice but not in male mice. CVB3 also increased the proportion and number of CD11a^hi^CD62L^lo^ CD8^+^ T cells in female mice, indicative of activation. Further, this response was independent of the inoculation route and type I interferon. Using a recombinant CVB3 virus expressing a model CD8^+^ T cell epitope, we found that the expansion of CD8^+^ T cells is viral-specific and not due to bystander activation. These data demonstrate that CVB3 induces a sex-dependent CD8^+^ T cell response and highlight the importance of sex-specific immune responses to viral pathogens.

## Introduction

Sex is a significant contributor to the outcome of various viral and bacterial infections. These sex-dependent outcomes are likely due to differences in the immune response between men and women. Previous studies have demonstrated that a heightened immune response in women promotes enhanced pathogen clearance compared to men (1). This heightened immune response is not without consequences, as females often experience more severe symptoms following infection and are more predisposed to autoimmune diseases. Moreover, sex-specific immunity also influence vaccine responses (2). Therefore, there is a critical need to understand the mechanism(s) that influence sex-dependent immune responses to pathogens.

T cells play an essential role in the host immune response to viral pathogens, and memory T cells contribute to vaccine-induced immunity (3-5). Several studies have demonstrated differences in the number and proportion of T cells in men and women. Women tend to have higher CD4+ T cell counts and a higher CD4/CD8 ratio than men (6). Activation of T cells is also enhanced in women. Upon PMA-ionomycin stimulation, female CD4+ T cells exhibit increased upregulation of TNFα, IFN-γ, and IL-17 (7). Similarly, activated female CD8+ T cells show enhanced upregulation of antiviral and proinflammatory genes compared to their male counterparts (8). Overall, these data indicate that sex differences in the response of T cells play a crucial role in the outcome of intracellular infections. However, questions remain about how sex differences in the T cell response impact specific viral infections.

Coxsackievirus B3 (CVB3) is a small, non-enveloped RNA virus in the Picornaviridae family. In humans, CVB3 is a primary cause of viral myocarditis. A sex bias in viral myocarditis has been observed where males are twice as likely to have severe sequela compared to females. This sex bias is also observed in the murine models of CVB3 (9, 10). In mice, the immune response to CVB3 has been implicated as a critical contributor to viral myocarditis. Acute CVB3-induced myocarditis is characterized by inflammatory infiltration of immune cells, including CD4^+^ and CD8^+^ T cells (11, 12). Several studies have shown that these T cells contribute to CVB3 clearance but can also facilitate disease following infection (13). Differences in the Th1 and Th2 CD4^+^ T cell responses between males and females influence myocarditis in mice (14, 15). Further, γδ T cells and sex hormones can tip the balance of this CD4^+^ T cell response to either promote or limit disease (16-19). Similarly, CD8^+^ T cells have also been shown to play dual roles in myocarditis. Depleting CD8^+^ T cells leads to increased viral titers in mice; however, CVB3 infection of CD8^+^-deficient, beta-2 microglobulin knockout mice are more resistant to the development of myocarditis (13). Unfortunately, efforts to elucidate CVB3-specific CD8^+^ T cell response have been hampered due to the limited ability to induce CD8^+^ T cell expansion *in vivo* (13-15, 20-23). Thus, the role of CD8^+^ T cells in CVB3 pathogenesis is still unclear.

CVB3 is a member of the enterovirus group that is transmitted through the fecal-oral route and initiates infection in the gastrointestinal tract. Previously we established an oral inoculation mouse model for CVB3 to recapitulate a natural route of infection through the intestine (24). This model uses C57BL/6 *Ifnar*^*-/-*^ (deficient for interferon α/β receptor) mice to facilitate viral replication in the intestine (24, 25). We recently determined that testosterone, the primary sex hormone in males, can promote intestinal CVB3 replication and viral dissemination in male mice (26). Further, gonadectomy of male mice to deplete endogenous testosterone completely protected males from CVB3-induced lethality. Similarly, testosterone can promote replication and dissemination in female mice, but these hormones are not associated with mortality in female mice. Further, ovariectomy of female mice does not alter the susceptibility to CVB3-induced mortality (24). Therefore, these data suggest sex hormones contribute to intestinal viral replication dissemination, but other unknown immune system components protect female mice from CVB3-associated lethality.

In this study, we sought to evaluate the CVB3-specific immune response in orally inoculated male and female mice. We found that CVB3 elicited a significant expansion of CD4^+^ and CD8^+^ T cells in female mice but not in male mice. We further evaluated the CD4^+^ and CD8^+^ T cell response in female mice for activation and found that female CD8^+^ T cells also had markers of activated CD8^+^ T cells. Further, we found that the expansion of female CD8^+^ T cells is independent of the route of inoculation and type I IFN. Finally, using a recombinant CVB3 strain (rCVB3-GP_33_) that has LCMV-specific CD8^+^ cytotoxic T lymphocyte GP33-41 epitope integrated into its polyprotein (20, 21, 27) we found that CD8^+^ T cell expansion was driven by viral antigen rather than bystander activation. These findings indicate a novel, sex-dependent CD8^+^ T cell response to CVB3 and further emphasize the inclusion of sex as a variable in the immune response to viral pathogens.

## Materials and Methods

### Cells and virus

HeLa cells were grown in Dulbecco’s modified Eagle’s medium (DMEM) supplemented with 10% calf serum and 1% penicillin-streptomycin at 37°C with 5% CO_2_. The CVB3-Nancy and CVB3-H3 infectious clones were obtained from Marco Vignuzzi (Pasteur Institute, Paris, France) and propagated in HeLa cells as described previously (24). CVB3 was quantified by a standard plaque assay using HeLa cells. The rCVB3-GP_33_ (rCVB3.6) was kindly provided to us by Lindsay Whitton and Taishi Kimura (Scripps Research Institute, La Jolla, California).

### Mouse experiments

All animals were handled according to the Guide for the Care of Laboratory Animals of the National Institutes of Health. All mouse studies were performed at Indiana University School of Medicine using protocols approved by the local Institutional Animal Care and Use Committee in a manner designated to minimalize pain, and any animals that exhibited severe disease were euthanized immediately by CO_2_ inhalation. C57BL/6 *PVR*^*+/+*^ wild-type and *Ifnar*^*-/-*^ mice were obtained from S. Koike (Tokyo, Japan) and were 8-14 weeks old at the time of infection (28). Mice were orally inoculated with 5×10^7^ PFU CVB3-Nancy in a final volume of 20 uL. For intraperitoneally injections, mice were inoculated with 1×10^4^ PFU of CVB3 or 1×10^8^ PFU of rCVB3.6 in a final volume of 200 uL of Phosphate Buffered Saline (PBS).

### Flow cytometry analysis

The spleen was collected from male and female mice at indicated time points and mechanically disrupted to generate a single-cell suspension. The erythrocytes were lysed using 1xRBC lysis buffer (BioLegend, catalog # 420301) and cells were then washed and incubated with TruStain fcX (CD16/CD32. Clone 93, BioLegend, catalog # 101320), and stained with surface antibodies for indicated immune cells. Next, cells were fixed using IC fixation buffer (Life Technologies (Fisher), catalog # 00-8222-49) and analyzed on a BD LSRFortessa flow cytometer and FlowJo software (BD Biosciences). The following mouse antibodies were used in an appropriate combination of fluorochromes: CD4 (clone GK1.5, BioLegend, catalog #100412), CD8a (clone 53-6.7, BioLegend, catalog #100725, #100711), MHC II (clone M5/114.15.2, BioLegend, catalog #107619), CD11a (clone N418, BioLegend, catalog #117327, 101124), CD19 (clone 6D5, BioLegend, catalog #115507) (1D3/CD19, BioLegend, catalog # 152404) NK1.1 (clone PK136, BioLegend, catalog # 108706) CD11c (clone N418, BioLegend, catalog # 117327), CD40 (eBioscience (Fisher), catalog # 17040182), CD80 (clone 16-10A1, BioLegend, catalog # 104707) CD86 (clone GL-1, BioLegend, 105013), CD62L (clone MEL-14, BioLegend, catalog # 104438), and CD49d (clone R1-2, BioLegend, catalog # 103607) GP33 (Ask Stephanie), CD3e (145-2C11, BioLegend, catalog # 100306) CD69 (clone H1.2F3, BioLegend, catalog # 104521), ly-6C (clone HK1.4, BioLegend, catalog # 128015) Ly-6G (clone 1A8, BioLegend, catalog # 127621), CD11b (clone M1/70, BioLegend, catalog # 101223) F4/80 (clone BM8, BioLegend, catalog # 123113). A representative gating strategy is provided in Supplementary Fig. 6.

### GP33 tetramer Staining

The H2D^b^ GP_33_ tetramer was provided as PE-conjugated by the NIH Tetramer Core facility. Spleens were collected and mechanically disrupted at 5, 8, and 15dpi to generate single-cell suspensions. Erythrocytes were lysed using 1xRBC lysis buffer, and then cells were incubated with TruStain fcX (anti-mouse CD16/CD32. Clone 93, BioLegend) for 15 mins at 1:50 dilution at 4 C and then incubated with the GP_33_ tetramer for 45 mins at 4 C at a final dilution of 1:400. Cells were washed and stained with surface antibodies for indicated immune cells.

### Statistical Analysis

Comparisons between control and study groups were analyzed using either an unpaired t-test or a one-way analysis of variance (ANOVA). Error bars in the figures represent the standard errors of the means. A p-value <0.05 was considered significant. All analyses were performed using GraphPad Prism 9 (GraphPad Software, La Jolla, CA).

## Results

### Expansion of CD4^+^ and CD8^+^ T cells occurs in female mice but not in male mice following oral inoculation with CVB3

We previously found that female *Ifnar*^*-/-*^ mice are protected against CVB3-induced lethality, and this protection did not correlate to intestinal replication and dissemination (26). Therefore, we hypothesized that sex-specific differences in immune responses might offer females protection against CVB3 following oral inoculation. To determine immune correlates of protection, we orally inoculated male and female C57BL/6 *Ifnar*^*-/-*^ mice with 5×10^7^ PFUs of CVB3-Nancy, collected the spleens at 5 days post-inoculation (dpi), and analyzed splenocytes by flow cytometry. We chose to examine immune cells at 5 dpi because we have previously shown that mortality in male mice typically begins at this time point (24, 29). At 5dpi, while we observed a significant increase in the numbers of splenic neutrophils and macrophages between uninfected male and female mice, we found no difference in the frequency and numbers of B cells, neutrophils, macrophages, and dendritic cells between uninfected and infected mice of either sex (Supplemental Figure 1). Contrary to other immune cells, we observed a significant decrease in the frequency of CD4^+^ T cells in infected male mice (Fig. 1A and 1B). The overall number of splenic CD4 T cells in CVB3-infected male mice was also reduced compared to uninfected male mice; however, this did not reach statistical significance (Fig 1C). Similarly, the frequency and number of CD8^+^ T cells in infected male mice were reduced compared to uninfected male mice, but this was not statistically significant (Fig. 1D and E). In female mice, we found no significant increase in the frequency of CD4^+^ and CD8^+^ T cells in infected or uninfected groups. However, in contrast to males, we found a significant increase in the number of splenic CD4^+^ and CD8^+^ T cells in CVB3-infected female mice compared to uninfected female mice (Fig. 1C and 1E). Moreover, the number of CD4^+^ and CD8^+^ T cells was significantly higher in CVB3-infected female mice compared to CVB3-infected male mice (Fig. 1C and 1E). Overall, these data indicate that female mice have increased numbers of splenic T cells following CVB3 infection.

**Figure 1.**
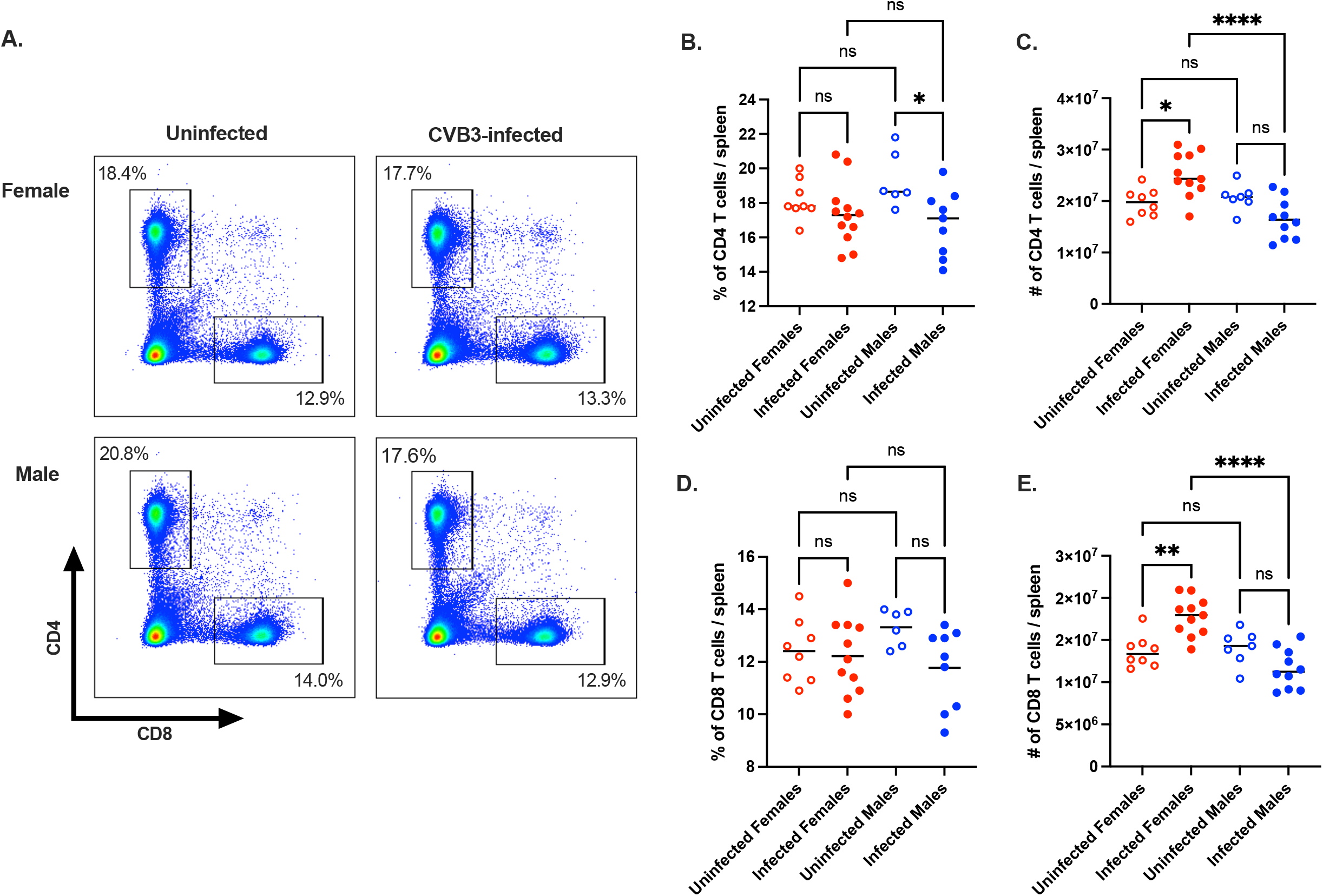
The CD4^+^ and CD8^+^ T cell response in CVB3-infected male and female *Ifnar*^*-/-*^ mice. Male and female *Ifnar*^*-/-*^ mice were orally inoculated with 5×10^7^ PFUs of CVB3. The spleen was harvested at 5dpi, and splenocytes were processed for analysis by flow cytometry. (A) Representative flow cytometry plots of T cells gated on CD4^+^ and CD8^+^ T cell expression in uninfected and infected male and female mice. The frequency (B) and number (C) of splenic CD4^+^ T cells 5dpi. The frequency (D) and number (C) of splenic CD8^+^ T cells 5dpi. Data points in the scatter plots represent individual mice, with lines representing the mean from two experiments. *p<0.05, **p<0.01, ****p<0.0001, One-way ANOVA.

### Activation and expansion of antigen-experienced CD8^+^ T cells occur in CVB3-infected female mice but not in male mice

Based on the increase in the numbers of CD4^+^ and CD8^+^ T cells in infected female mice, we hypothesized that these T cells were activated in female mice following CVB3 infection. First, we assessed CD8^+^ T cell activation by the expression of CD62L. Naïve CD8^+^ T cells are CD62L^hi^, and activated CD8^+^ T cells differentiate into effector subtypes during acute infections. During this effector phase, CD62L is downregulated (30-32). Following CVB3 inoculation, we observed a significant increase in the frequency and number of CD62L^lo^ CD8^+^ T cells in infected female mice compared to uninfected female mice (Fig. 2A – 2C). Next, we investigated the expression of the integrin molecule CD11a on CD62L^lo^ CD8^+^ T cells. Upregulation of CD11a can distinguish naïve CD8^+^ T cells from effector and memory CD8^+^ T cells (30). We found that infection with CVB3 significantly increased the frequency and number of CD11a^hi^CD62L^lo^ CD8^+^ T cells in female mice (Fig. 2D – 2F). Taken together, these data indicate that CVB3 alters the frequency and number of CD62L^lo^ and CD11a^hi^CD62L^lo^ CD8^+^ T cells in female mice, indicative of CD8^+^ T cell activation.

**Figure 2.**
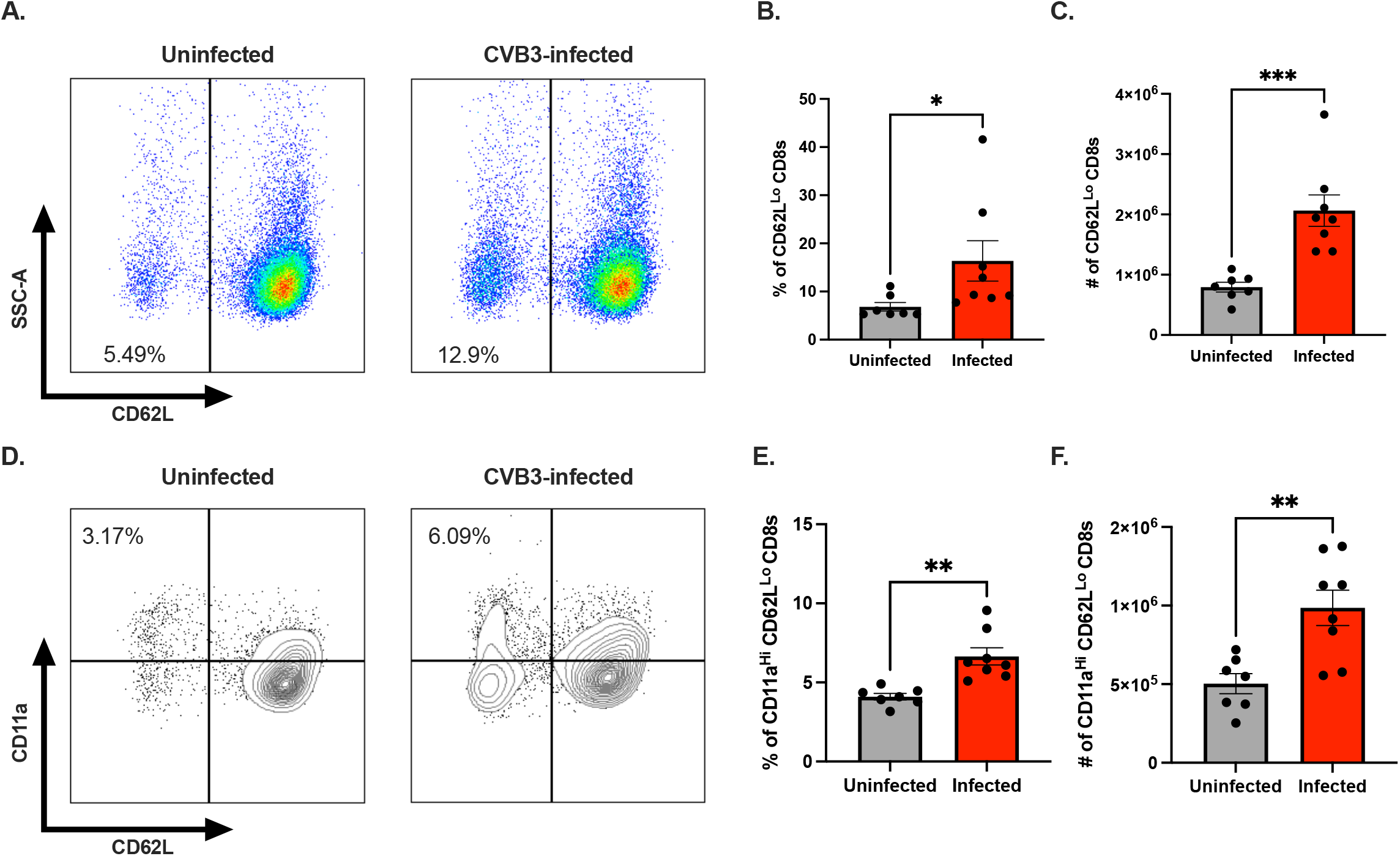
CVB3 induces an expansion of activated CD8^+^ T cells in female *Ifnar*^*-/-*^ mice. (A) Representative flow cytometry plots of CD62L expression on CD8^+^ T cells in uninfected and infected female *Ifnar*^*-/-*^ mice. The frequency (B) and number (C) of CD62L^lo^ CD8^+^ T cells in the spleen 5dpi in *Ifnar*^*-/-*^ mice. (D) Representative flow cytometry plots of CD11a and CD62L expression gated on CD8^+^ T cells in uninfected and infected female *Ifnar*^*-/-*^ mice. The frequency (E) and number (F) of CD11a^hi^CD62L^lo^ CD8^+^ T cells in female mice 5dpi. All data are from two independent experiments with n = 7-8 mice per group and are shown as mean ± SEM. *p<0.05, **p<0.01, ***p<0.001, unpaired t-test.

The loss of CD62L on CD8^+^ T cells can occur through bystander activation rather than direct antigen-specific engagement of the T cell receptor. Since we observed an increase in CD62L^lo^ CD8 T cells, we sought to determine if the activation of CD8 T cells in female mice was due to encountering CVB3 antigen or bystander activation. However, identifying if CVB3 generates a viral-specific CD8^+^ T cell response requires known immunodominant T cell epitopes. Unfortunately, CVB3-specific CD8^+^ T cell epitopes have not been clearly defined. To overcome this limitation, we examined antigen-experienced T cells using surrogate markers. Previous studies have established that antigen-experienced T cells induced following infection can be tracked, regardless of their specificity, using CD11a^hi^CD49d^+^, and CD8a^lo^CD11a^hi^ surrogate markers for CD4^+^ and CD8^+^ T cells, respectively (30, 33-37). To examine antigen-experienced CD8^+^ T cells, we analyzed the frequency and number of CD8a^lo^CD11a^hi^ CD8^+^ T cells in the spleen at 5dpi (Fig. 3A). We found that the frequency of antigen-experienced CD8a^lo^CD11a^hi^ CD8^+^ T cells was not significantly different between infected and uninfected mice of either sex at 5dpi (Fig. 3B). In contrast, while we did not see any difference in the number of CD8a^lo^CD11a^hi^ CD8^+^ T cells between uninfected and infected male mice, we observed a significant increase in the number of CD8a^lo^CD11a^hi^ CD8^+^ T cells in infected female mice compared to uninfected female mice (Fig. 3C). These data indicate that CVB3 induces a viral-specific CD8 T cell response in female mice and this T cell response can be tracked using the surrogate-maker approach.

**Figure 3.**
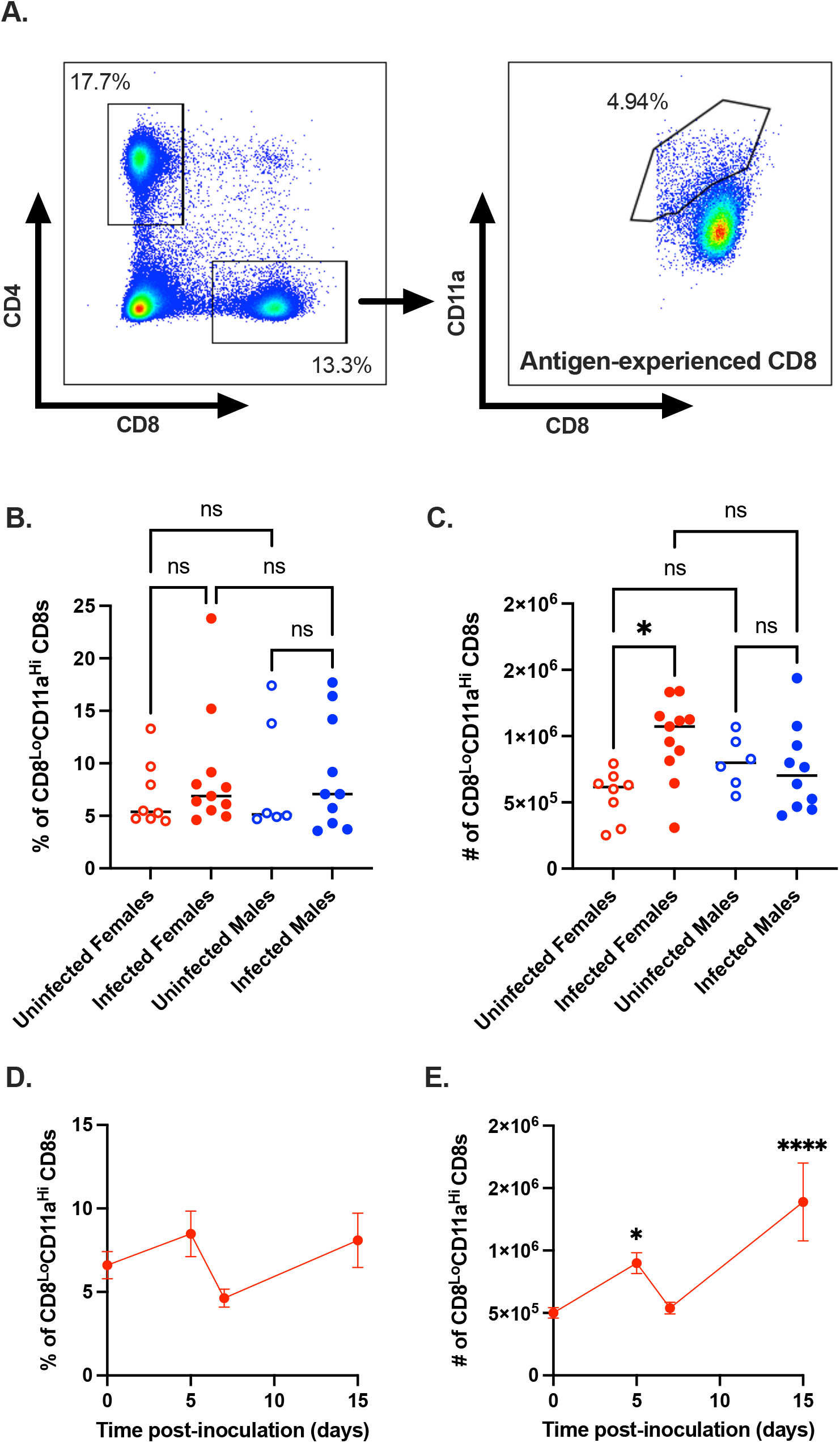
CVB3 induces expansion of antigen-experienced CD8^+^ T cells. (A) Representative flow cytometry plots for the gating strategy of CD8a^lo^CD11a^hi^ expression on CD8^+^ T cells. The frequency (B) and number (C) of CD8a^lo^CD11a^hi^ CD8^+^ T cells in male and female *Ifnar*^*-/-*^ mice 5dpi. Data points in the scatter plots represent individual mice, with lines representing the mean from two experiments. *p<0.05, One-way ANOVA. The kinetics of the frequency (D) and number (E) of CD8a^lo^CD11a^hi^ CD8^+^ T cells in female *Ifnar*^*-/-*^ mice from the spleen. All data are from two-three independent experiments with n = 7-14 mice per time point and are shown as mean ± SEM. *p<0.05, ****p<0.0001, unpaired t-test.

Finally, to examine the kinetics of the antigen-experienced CD8 T cell response in female mice, we orally inoculated female *Ifnar*^*-/-*^ mice with CVB3 and examined the total CD8a^lo^CD11a^hi^ CD8^+^ T cell response at 5, 7, and 15dpi. We found that while the proportion of total CD8a^lo^CD11a^hi^ CD8^+^ T cells did not change over time in response to CVB3 (Fig. 3D), there were two peaks in the increase in the number of splenic CD8a^lo^CD11a^hi^ CD8^+^ T cells. At 5 and 15dpi, we found CVB3 induced a significant increase in the splenic numbers of CD8a^lo^CD11a^hi^ CD8^+^ T cells compared to uninfected female mice (Fig. 3E). Further, there was a contraction in the number of CD8a^lo^CD11a^hi^ CD8^+^ T cells on day 7 post-infection. Taken together, our data indicate that CVB3 induces activation and expansion of CD8^+^ T cells at 5 and 15dpi in female mice following CVB3 infection.

### Antigen-experienced CD11a^hi^CD49d^+^ CD4^+^ T cells expand in female mice following CVB3 infection

CD4^+^ T cells have been previously shown to play a significant role in the CVB3-induced myocarditis (14, 15, 23, 38). To further characterize the T cell response in male and female *Ifnar*^*-/-*^ mice, we examined the response of CVB3-specific CD4^+^ T cells using a similar surrogate marker approach (30, 33-37). Following oral inoculation, we observed no significant difference in the frequency and numbers of CD11a^hi^CD49d^+^ CD4^+^ T cells at 5dpi in infected mice versus uninfected mice of either sex (Supplemental Fig 2A - C). However, the numbers of CD11a^hi^CD49d^+^ CD4^+^ T cells were significantly higher in infected female mice compared to infected male mice (Supplemental Fig 2C). These data suggest that CVB3 infection may induce a minimal antigen-experienced CD4^+^ T cell response at 5dpi, which is higher in females than males. Next, we analyzed the CD11a^hi^CD49d^+^ CD4^+^ T cell response in female mice over time following CVB3 infection. In contrast to antigen-experienced CD8^+^ T cells, we found that the frequency and number of CD11a^hi^CD49d^+^ CD4^+^ T cells peaked at 15dpi (Supplemental Fig 2D and 2E). Thus, these data indicate that CVB3 induces the expansion of antigen-experienced CD4^+^ T cells in female mice; however, this expansion occurs after the initial expansion of CD8^+^ T cells.

### Bypassing the initial CVB3 infection through the intestinal mucosa does not impact the CD8^+^ T cell response in female mice

Previous studies showing a limited CD8^+^ T cell response in mice were based on a systemic infection model using intraperitoneal (ip) injections as the route of inoculation. Since the intestinal mucosa can promote robust immunity, we hypothesized that the previous lack of an observed CD8^+^ T cell response might be due to a systemic infection that bypasses the initial infection of the intestine. To investigate this hypothesis, we ip inoculated male and female *Ifnar*^*-/-*^ mice with 1×10^4^ PFUs of CVB3 to bypass the intestine. At 5dpi, the spleens were harvested, and splenocytes were analyzed by flow cytometry. Similar to our oral inoculation model, we observed no significant difference in the numbers of B cells, macrophages, neutrophils, and dendritic cells between infected and uninfected male and female mice (Supplemental Figure 3). Further, we found that CVB3 infection leads to a significant decrease in the frequency of CD4^+^ T cells in males and female mice, but only a significant decrease in the frequency of CD8^+^ T cells in males (Table 1). In contrast to our hypothesis, we found that CVB3 increases the proportion and number of antigen-experienced CD8a^lo^CD11a^hi^ CD8^+^ T cells in infected female mice compared to uninfected female mice, similar to our findings from oral inoculation (Fig. 4A and 4B). CVB3 also increased the frequency of antigen-experienced CD8^+^ T cells in male mice; however, no significant difference was observed in the number of CD8a^lo^CD11a^hi^ CD8^+^ T cells between infected or uninfected male mice (Fig. 4A and 4C). These data suggest that the CD8^+^ T cell response in female mice is independent of the inoculation route.

**Table 1.**

| | Cell type | Frequency of cells (%) per spleen | | Total number of cells ($\times 10^4$) per spleen | |
| --- | --- | --- | --- | --- | --- |
|  |  | Uninfected | Infected | Uninfected | Infected |
| **Male** | CD4 T cells | 19.49±0.69* | 15.23±0.54* | 524.91±52.98 | 399.04±29.64 |
| | CD8 T cells | 13.98±0.44* | 11.66±0.44* | 393.22±40.53 | 312.98±23.49 |
| **Female** | CD4 T cells | 19.44±0.71* | 16.75±0.51* | 501.56±38.81 | 540.85±70.99 |
| | CD8 T cells | 13.78±0.67 | 12.14±0.34 | 344.92±30.53 | 376.51±46.94 |

Data are presented as mean ± SEM from two independent experiments (n=13-16 per group).
Significant differences between groups are denoted by asterisks (One-way ANOVA, \*p<0.05).

**Figure 4.**
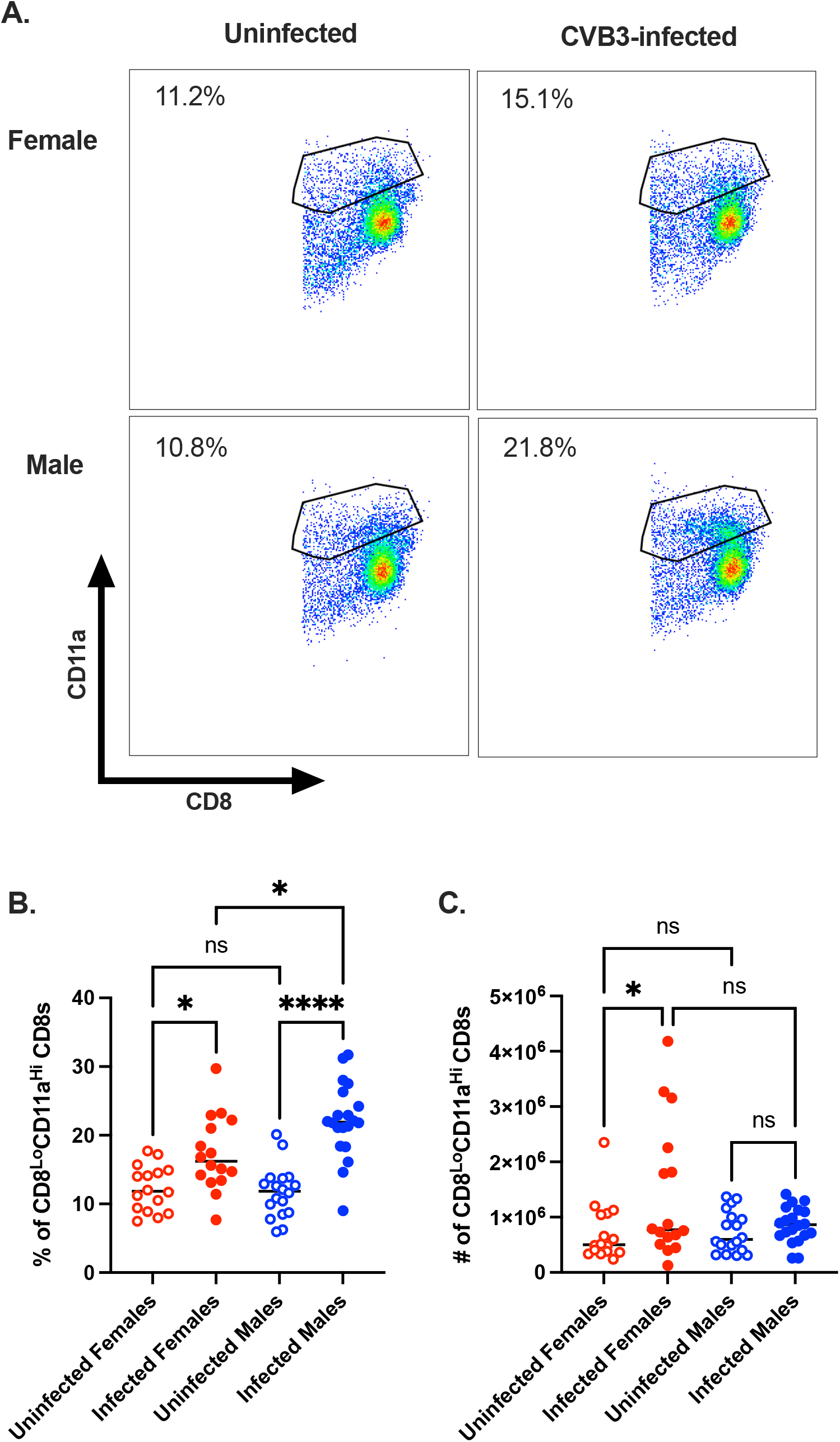
Bypassing infection through the oral mucosa does not impact the expansion of antigen-experienced CD8^+^ T cells in female mice. Male and female *Ifnar*^*-/-*^ mice were ip inoculated with 1×10^4^ PFU of CVB3. The spleen was harvested at 5dpi, and splenocytes were analyzed by flow cytometry. (A) Representative flow cytometry plots of CD8a^lo^CD11a^hi^ expression on CD8^+^ T cells from uninfected and infected male and female *Ifnar*^*-/-*^ mice following ip inoculation with CVB3. The frequency (B) and number (C) of CD8a^lo^CD11a^hi^ CD8^+^ T cells in uninfected and infected male and female *Ifnar*^*-/-*^ mice 5dpi. Data points in the scatter plots represent individual mice, with lines representing the mean from three experiments. *p<0.05, ****p<0.0001, One-way ANOVA.

Next, to examine the CD4^+^ T cell response, we examined antigen-experienced CD11a^hi^CD49d^+^ CD4^+^ T cells. Similar to our findings in *Ifnar*^*-/-*^ mice, we found no statistical differences in the frequency of CD11a^hi^CD49d^+^ CD4^+^ T cells between uninfected and infected mice of both sexes at 5 dpi (Supplemental Fig. 4A and B). In contrast, while we observed no difference in the number of CD11a^hi^CD49d^+^ CD4^+^ T cells in infected and uninfected female mice, we observed a significant decrease in the number of CD11a^hi^CD49d^+^ CD4^+^ T cells in infected male mice compared to uninfected male mice (Supplemental Fig. 4C). Taken together, these data indicate that bypassing the intestinal mucosa and infecting mice via ip injection still generates an expansion of antigen-experienced CD8^+^ T cells in female *Ifnar*^*-/-*^ mice while having little impact on antigen-experienced CD4^+^ T cells in both sexes.

### The type I interferon response does not impact the CD8^+^ T cell response in female mice

Inflammatory cytokines, including type I interferons (IFN), have been previously shown to be necessary for the activation, proliferation, and differentiation of CD8^+^ T cells (3). Since our data show T cell expansion in female mice that lack the type I IFN receptor, we examined the CD8^+^ T cell response in female wild-type C57BL/6 mice to determine the impact of type I IFN on this response. Following ip inoculation, we observed no difference in the frequency of splenic CD8^+^ T cells between uninfected and infected wild-type female mice (Fig. 5A). However, the total numbers of splenic CD8^+^ T cells trended lower in infected female mice that reached near significance (p=0.0599) (Fig. 5B). In contrast, infected female wild-type mice had a higher frequency of CD8a^lo^CD11a^hi^ CD8^+^ T cells compared to uninfected female mice (Fig. 5C). The overall number of CD8a^lo^CD11a^hi^ CD8^+^ T cells remained similar between uninfected and infected female mice (Fig. 5D). However, this is likely due to the reduction in CD8 T cells in infected female mice. Taken together, these data indicate that CVB3 leads to an increased proportion of antigen-experienced CD8^+^ T cells in wild-type female mice, and the loss of type I IFN does not impact this CD8^+^ T cell response.

**Figure 5.**
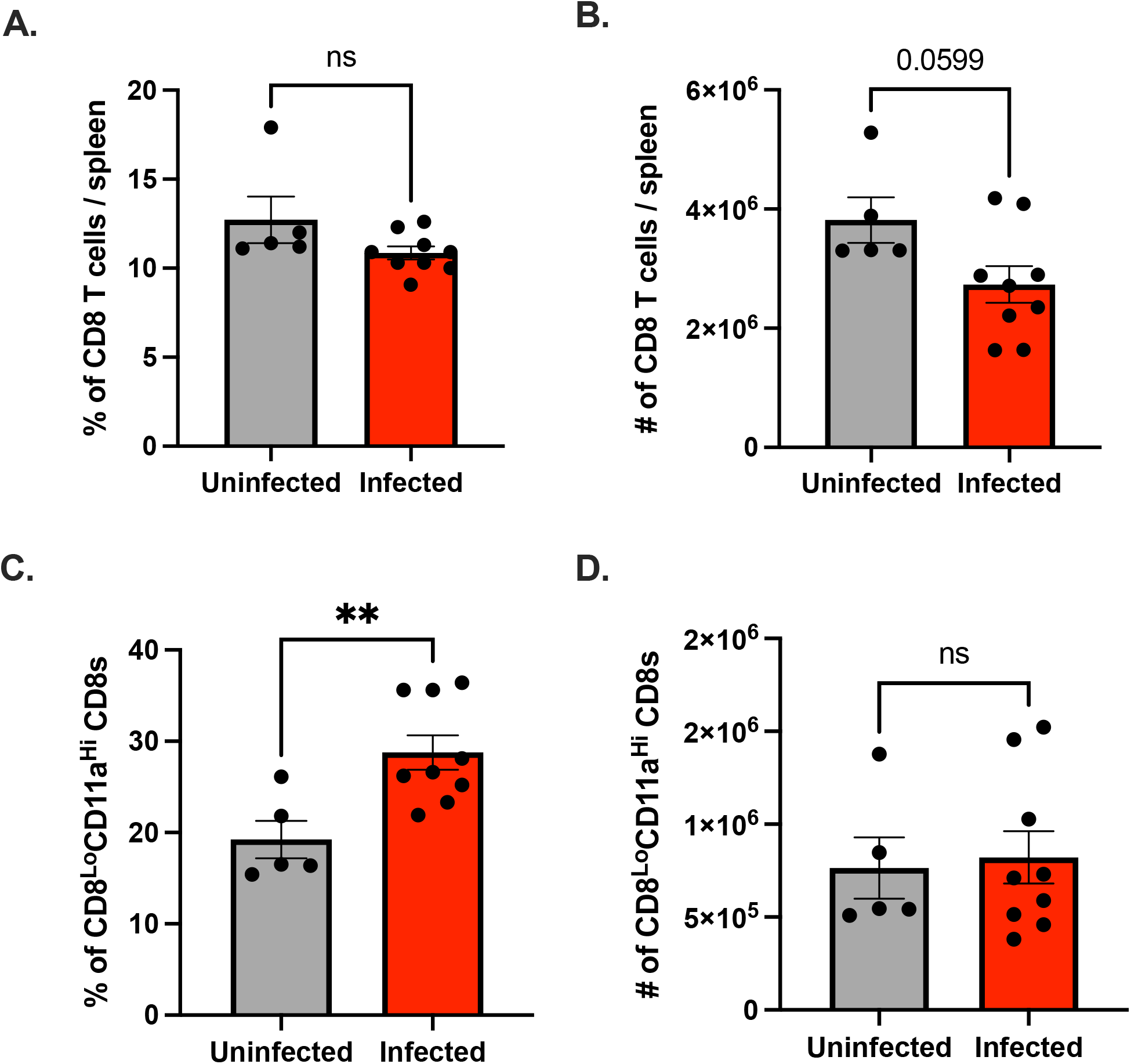
Type I IFN does not impact the expansion of antigen-experienced CD8^+^ T cells in female mice. Wild-type female C57BL/6 mice were ip inoculated with 1×10^4^ PFUs of CVB3. The spleen was harvested at dpi, and splenocytes were processed for analysis by flow cytometry. The frequency (A) and number (B) of CD8^+^ T cells in the spleen of wild-type female C57BL/6 mice 5dpi. The frequency (C) and number (D) of CD8a^lo^CD11a^hi^ CD8^+^ T cells in wild-type female C57BL/6 mice 5dpi. All data are from two independent experiments and are shown as mean ± SEM. **p<0.01, unpaired t-test.

### Antigen-experienced CD8 T cells expand due to CVB3 antigen and not due to bystander activation

Previous studies have shown that CVB3 fails to trigger viral-specific CD8^+^ T cells *in vivo*; however, these studies have primarily been carried out in male mice (20-22). Since we observed an expansion of CD8a^lo^CD11a^hi^ CD8^+^ T cells in female mice, we evaluated if this CD8 T cell response was viral-specific or represented the expansion of bystander CD8^+^ T cells. To address this question, we used a recombinant CVB3 (rCVB3-GP_33_) that encodes a well-characterized CD8 T cell epitope from lymphocytic choriomeningitis virus (LCMV) (Fig. 6A). This recombinant CVB3, while attenuated *in vivo*, still leads to a productive infection, generating high tissue titers, and is cleared similarly to wtCVB3 (20-22). We ip inoculated female *Ifnar*^*-/-*^ mice with 10^8^ PFU of rCVB3-GP_33_ or 10^4^ PFU of wild-type CVB3 (wtCVB3) as a control. As a control for gating GP_33_-specific CD8^+^ T cells, we also infected female *Ifnar*^*-/-*^ mice with LCMV (Supplemental figure 5). The spleen was harvested from infected mice at 15dpi, and virus-specific CD8^+^ T cells were analyzed by flow cytometry using an H2-D^b^ GP_33_ tetramer (Fig. 6B). At 15dpi, we observed a significant increase in the frequency and number of GP_33_-specific CD8 T cells in the spleen from female mice infected with the rCVB3-GP_33_ virus compared to female mice infected with wtCVB3 (Fig. 6C and 6D). Overall, these data indicate that CVB3 drives the expansion of virus-specific CD8^+^ T cells in female mice.

**Figure 6.**
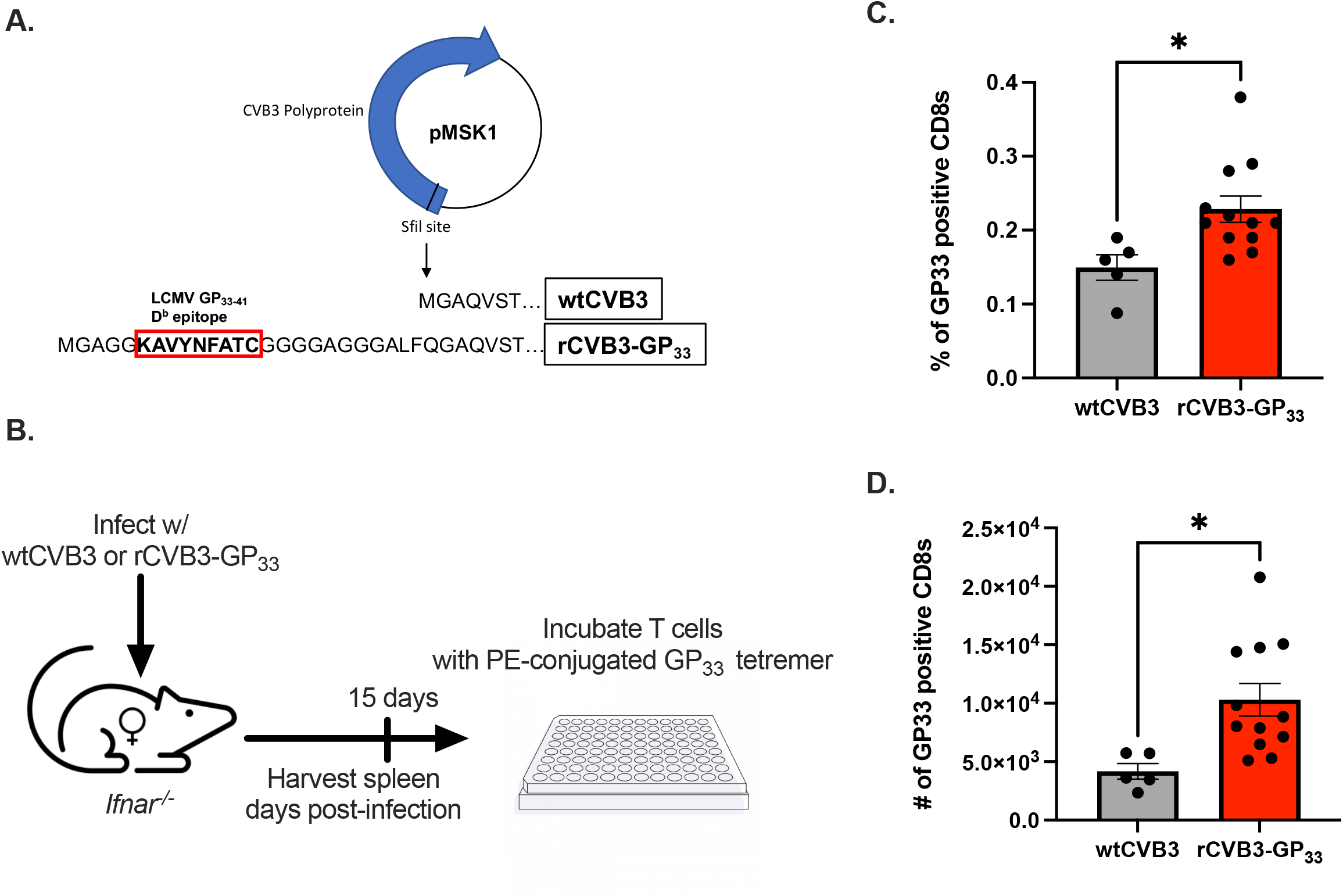
CVB3 induces a virus-specific expansion of CD8^+^ T cells in female mice. (A) the rCVB3-GP_33_ amino acid sequence encoding the LCMV GP33-41 CD8^+^ T cell epitope (adapted from (20)). (B) Schematic of the experimental design to determine the virus-specific CD8^+^ T cell response in female *Ifnar*^*-/-*^ mice. The frequency (C) and number (D) of CD8^+^ T cells staining with H2-D^b^ GP_33_ tetramer in female *Ifnar*^*-/-*^ mice 15dpi. All data are from two independent experiments and are shown as mean ± SEM. *p<0.05, unpaired t-test.

## Discussion

Numerous studies have shown that sex is a significant variable in infectious diseases, and the immune response has been associated with this sexual dimorphism (1, 6, 39). Similarly, sex-specific immune responses have been characterized in CVB3-induced myocarditis in mice. In this study, we examined the immune response in male and female mice following oral inoculation to identify potential immune correlates of protection. Our data indicate that female, but not male, CD8^+^ T cells are activated and respond to CVB3-specific antigen following infection. Overall, these data suggest that the CD8^+^ T cell response represents another sex-dependent immune factor during CVB3 infections.

Previous studies have indicated that CVB3 fails to elicit a robust CD8^+^ T cell response. In humans and mice, few CVB3-specific CD8^+^ T cell epitopes have been identified, and the frequency of viral-specific CD8^+^ T cells is extremely low in response to infection (40-44). Further, using a recombinant CVB3 that expresses the GP33 immunodominant peptide from LCMV, minimal differences in the CD8^+^ T cell activation markers, CD62L and CD11a, were detected. The use of a GP33 tetramer also failed to identify any CVB3-specific CD8^+^ T cells following infection (21). However, these conclusions were primarily drawn based on data from the infection of male mice. Our data confirm these previous findings as we found no significant indication of a detectable CD8^+^ T cell response in male mice. We observed a trending decrease in the frequency and number of male CD8^+^ T cells following infection in *Ifnar*^*-/-*^ mice (Fig. 1D and 1E). Further, the proportion and number of antigen-experienced CD8a^lo^CD11a^hi^ CD8^+^ T cells were similar between uninfected and infected males (Fig. 3B and 3C). One caveat we observed is that the proportion of antigen-experienced CD8^+^ T cells following ip inoculation of *Ifnar*^*-/-*^ males is significantly higher in infected male mice compared to uninfected male mice. However, this increase was not seen in the overall numbers, suggesting that the trending decrease in overall CD8^+^ T cells in males was the reason for the overall increase in the frequency of these CD8^+^ T cells. In contrast to males and the previous studies, our data suggest that CD8^+^ T cells expand in CVB3-infected female mice. Using a surrogate marker approach and a recombinant CVB3 expressing a well-characterized CD8^+^ T cell epitope, we found significant increases in the numbers of antigen-experienced CD8^+^ T cells (Fig. 3 and Fig. 4). Further, T cells in female mice had markers of activation (Figure 2A-G). Therefore, our data indicate a sex-specific CD8^+^ T cell response to CVB3. While we did not investigate it in this study, we speculate that CVB3 elicits female CD8^+^ T cells to differentiate into effector subtypes. Examining this differentiation will be the focus of future studies.

What might explain the sex differences in the expansion of CD8^+^ T cells to CVB3? A few possibilities exist; first, females may have an enhanced ability to prime CD8^+^ T cells in response to CVB3. Previous studies have shown that CVB3 inhibits antigen presentation by limiting the surface expression of MHC I in male dendritic cells (22). However, antigen-presenting cells (APCs) in female mice are reported to upregulate more MHC molecules in response to stimulus than in male mice (45). Further, the pattern-recognition receptor, toll-like receptor 7 (TLR7), is located on the X chromosome. Female dendritic cells express higher levels of TLR7, which can lead to an increased interferon response (46-48). Since the ligand for TLR7 is single-stranded RNA that recognizes ssRNA viruses such as Coxsackievirus B3, female APCs may be inherently better at detecting and priming T cells in response to CVB3. We speculate that the ability of CVB3 to limit the expression of co-stimulatory molecules and MHC I may be restricted to males, and females may overcome this inhibition. Further studies are required to dissect the sex-specific activation and antigen presentation of dendritic cells following CVB3 infection.

Another possibility for this sex difference may be the cytokine signals required to activate CD8^+^ T cells. Type I IFNs have been identified as an important signal for CD8^+^ T cells, and the lack of type I IFN signaling can severely blunt the viral CD8^+^ T cell response (49). Further, type I IFNs also provide a survival signal for CD8^+^ T cells during a clonal expansion (49, 50). Our data indicate that the expansion of female CD8^+^ T cells is independent of type I IFN, since we observed a similar expansion of female antigen-experienced CD8^+^ T cells in both wild-type and *Ifnar*^*-/-*^ (Fig. 3 and 4). However, the impact of type I IFN on CD8^+^ T cell survival may explain the kinetics of CD8a^lo^CD11a^hi^ CD8^+^ T cells in CVB3-infected female mice. We observed an initial increase in the numbers of CD8a^lo^CD11a^hi^ CD8^+^ T cells in female *Ifnar*^*-/-*^ mice at 5dpi, and there was a subsequent contraction at 7dpi (Fig. 3E). This may be due to the lack of IFN stimulation required for T cell survival or continued expansion. Interestingly, the numbers of CD8a^lo^CD11a^hi^ CD8^+^ T cells rebound on day 15, coinciding with the increase in antigen-experienced CD4^+^ T cells. Thus, our data indicate that CVB3 can induce expansion of female CD8^+^ T cells without type I IFN signaling, which may or may not impact the survival of these cells. Further studies are required to examine the long-term effects of type I IFN on female CD8^+^ T cells and if these CD8^+^ T cells differentiate into effector and memory subtypes.

Since type I IFN does not impact early expansion, our data suggest that other potential cytokines may serve as the third signal required to activate and lead to the expansion of female CD8^+^ T cells. We speculate that two cytokines, interleukin-21 (IL-21) and interleukin-12 (IL-12), may serve in this role. IL-21 has been previously implicated in promoting the activation of CD8^+^ T cells during CVB3 infection. IL-21 receptor (IL-21R) knockout mice have lower CD8^+^ T cell counts following CVB3 infection. Further, infected mice with transplanted IL-21R CD8^+^ T cells have less CVB3-induced myocarditis than those transplanted with wild-type CD8^+^ T cells (51). Additionally, IL-12 has been shown to overcome the lack of type I IFN signaling on CD8^+^ T cell activation and expansion. *Ex vivo* peptide stimulation of type I IFN-receptor knockout CD8^+^ T cells in the presence of recombinant IL-12 restored clonal expansion to wild-type levels (49). Further, *in vitro* antigen stimulation of naïve CD8^+^ T cells with IL-12 stimulated expansion and effector differentiation, similar to type I IFN (52). Interestingly, IL-12 has also been shown to play a role in a sex-specific T cell response to *Listeria monocytogenes*. In response to bacterial infection, an enhanced capacity for female CD8^+^ T cells to respond to IL-12 led to a higher proportion of short-lived effector CD8^+^ T cells (53). In contrast, *L. monocytogenes* infection drove male CD8^+^ T cells toward memory precursor effector cells. Further work is required to identify if IL-21, IL-12, or other cytokine signals activate CD8^+^ T cells in a sex-dependent manner and how these cytokines drive the T cell response to CVB3.

Finally, sex hormones may directly or indirectly impact CD8^+^ T cell activation, expansion, and differentiation. While differences in the absolute numbers of CD8^+^ T cells have been identified in men and women, T cells express the classical sex hormone receptors. Our lab and others have shown that sex hormones contribute to CVB3 pathogenesis. Recently we found that castrated males, depleted of endogenous testosterone, have a higher proportion and number of CD11a^hi^CD62L^lo^ CD8^+^ T cells compared to testosterone-treated males following CVB3 infection (26). These data suggest that testosterone can impact CD8^+^ T cell activation in CVB3 infections.

How testosterone and other sex hormones such as estrogens and progesterone directly act on CVB3-specific CD8^+^ T cells is unclear. Still, we cannot exclude the possibility that these hormones may indirectly alter the T cell response through modulating the APC or the cytokine signals. Studies to evaluate the role of sex hormones more directly on the CD8^+^ T cell response are ongoing in our laboratory.

CVB3 is an enterovirus that initiates replication in the intestine. Recent data suggest that adenovirus and murine norovirus recruit Ly6A^+^CCR9^+^CD4^+^ T cells to the intestinal epithelium, which help control viral replication (54). However, this CD4^+^ T cell response was not observed with reovirus infection, suggesting this T cell response is viral-specific. If intestinal T cells can control CVB3 replication and viral dissemination is unclear; however, immune correlates of protection against CVB3 may lie outside the intestine. In agreement, testosterone treatment of female mice enhances intestinal CVB3 replication, dissemination, and increases viral loads in peripheral tissues, but testosterone-treated female mice are still protected from CVB3-induced mortality. Therefore, the peripheral immune response may be necessary for controlling lethality, which may be advantageous for vaccine design. Further, CVB3 generated a CD8^+^ T cell response in female mice regardless of the inoculation route, which may also influence vaccine design.

In summary, our data yield new insight into the regulation of the CD8^+^ T cell response to CVB3 and highlight the immune differences between males and females. Our findings demonstrate that the CD8^+^ T cell response to CVB3 is sex-dependent. Future studies are required to examine if female CD8^+^ T cells differentiate into effector subtypes and the ability of CVB3 to induce immune memory in these mice. Overall, these data further strengthen the idea that biological sex is an important variable that should be considered when evaluating immune responses to viral pathogens.

## Acknowledgments

We would like to thank the members of the Indiana University Melvin and Bren Simon Cancer Center Flow Cytometry Resource Facility for their outstanding technical support. We thank the NIH Tetramer Core Facility (contract number 75N93020D00005) for providing the GP33 tetramer.

## Footnote

- This work was funded by a K01 DK110216, R03 DK124749, and a Biomedical Research Grant from the Indiana Clinical and Translational Sciences Institute to CMR. The Indiana University Melvin and Bren Simon Comprehensive Cancer Center Flow Cytometry Resource Facility (FCRF) is funded in part by NIH, National Cancer Institute (NCI) grant P30 CA082709 and National Institute of Diabetes and Digestive and Kidney Diseases (NIDDK) grant U54 DK106846. The FCRF is supported in part by NIH instrumentation grant 1S10D012270.

## Abbreviations used in this article

CVB3: Coxsackievirus B3
Dpi: days post-inoculation
IFN: interferon
ip: intraperitoneal

## Figure Legends

**Supplemental Figure 1**. Splenic immune cell responses in *Ifnar*^*-/-*^ mice to CVB3 following oral inoculation. Male and female *Ifnar*^*-/-*^ mice were orally inoculated with 5×10^7^ PFUs of CVB3. The frequency and number of splenic CD19^+^ B cells (A, B), neutrophils (C, D), macrophages (E, F), and monocytes (G, H). ns, not significant. *p<0.5, **p<0.01, One-way ANOVA.

**Supplemental Figure 2**. CVB3 induces expansion of antigen-experienced CD4^+^ T cells. (A) Representative flow cytometry plots for the gating strategy of CD11a^hi^ CD49d^+^ expression on CD4^+^ T cells. The frequency (B) and number (C) of CD11a^hi^ CD49d^+^ CD4^+^ T cells in male and female *Ifnar*^*-/-*^ mice 5dpi. Data points in the scatter plots represent individual mice, with lines representing the mean from two experiments. **p<0.01, One-way ANOVA. The kinetics of the frequency (D) and number (E) of CD8a^lo^CD11a^hi^ CD8^+^ T cells in female *Ifnar*^*-/-*^ mice from the spleen. All data are from two-three independent experiments with n = 7-14 mice per time point and are shown as mean ± SEM. ****p<0.0001, unpaired t-test.

**Supplemental Figure 3**. Splenic immune cell responses in *Ifnar*^*-/-*^ mice to CVB3 following ip inoculation. Male and female *Ifnar*^*-/-*^ mice were ip inoculated with 1×10^4^ PFUs of CVB3. The frequency and number of splenic CD19^+^ B cells (A, B), neutrophils (C, D), macrophages (E, F), and monocytes (G, H). ns, not significant, One-way ANOVA.

**Supplemental Figure 4**. Antigen-experienced CD4^+^ T cells in CVB3 infected female mice inoculated by the ip route. Male and female *Ifnar*^*-/-*^ mice were ip inoculated with 1×10^4^ PFU of CVB3. The spleen was harvested at 5dpi, and splenocytes were analyzed by flow cytometry. (A) Representative flow cytometry plots of CD11a^hi^ CD49d^+^ expression on CD4^+^ T cells from uninfected and infected male and female *Ifnar*^*-/-*^ mice following ip inoculation with CVB3. The frequency (B) and number (C) of CD11a^hi^ CD49d^+^ CD4^+^ T cells in uninfected and infected male and female *Ifnar*^*-/-*^ mice 5dpi. Data points in the scatter plots represent individual mice, with lines representing the mean from three experiments. *p<0.05, One-way ANOVA.

**Supplemental Figure 5**. Identification of GP33-tetramer positive CD8^+^ T cells following LCMV infection. (A) Schematic of the experimental design. Female *Ifnar*^*-/-*^ mice were ip inoculated with 2×10^5^ PFU of LCMV, and at 8dpi, the spleen was harvested for flow cytometry analysis. (B) Representative flow cytometry plot for the gating strategy to identify GP33-tetramer positive CD8^+^ T cells in LCMV-infected female *Ifnar*^*-/-*^ mice.

**Supplemental Figure 6**. Representative gating strategies. Representative gating strategies to identify CD62L^lo^ CD8^+^ T cells, of CD11a^hi^CD62L^lo^ CD8^+^ T cells, and CD8a^lo^CD11a^hi^ CD8^+^ T cells.

## Notes

### Competing Interest Statement

The authors have declared no competing interest.

